# Age-dependent visualization of neural progenitor cells within the rostral migratory stream via MRI and endogenously labeled micron-sized iron oxide particles

**DOI:** 10.1101/429787

**Authors:** Dorela D. Shuboni-Mulligan, Shatadru Chakravarty, Christiane L. Mallett, Alexander M. Wolf, Stacey Forton, Erik M. Shapiro

## Abstract

The subventricular zone (SVZ) is one of the primary sources for rodent neural progenitor cells (NPC), however, aging greatly impacts the substructure of the region and rate of new cell birth. To determine if age impacts the rate of *in vivo* migration within animals, we examined the rostral migratory stream (RMS) of animals across 12 days using an established MRI technique. To visualize NPCs, we injected micron sized particles of iron oxide (MPIO) into the lateral ventricle to endogenously label cells within the SVZ, which then appeared as hypo-intensive spots within MR images. Our *in vivo* MRI data showed that the rate of migration was significantly different between all ages examined, with decreases in the distance traveled as age progressed. The total number of iron oxide labeled cells within the olfactory bulb on day 12, decrease significantly when compared across ages in *ex vivo* high-resolution scans. We also, for the first time, demonstrated the endogenous labeling of cells within the dentate gyrus (DG) of hippocampus. Here too, there was a significant decrease in the number of labeled cells within the structure across age. Histology of the NPCs verified the decrease in labeling of cells with doublecortin (DCX) as age progressed for both regions. The dramatic reduction of labeling in NPCs within the SVZ and DG observed with MRI, demonstrates the importance of understanding the impact of age on the relationship of NPC and disease.

## Introduction

Neurogenesis within the adult rodent brain primarily occurs in two regions: the subventricular zone (SVZ) of the lateral ventricle and the subgranular zone (SGZ) of the hippocampus (Altman, 1969; Kaplan & Hinds, 1977; Kaplan & Bell, 1984). Neural stem cells within the SVZ give rise to neural progenitor cells (NPC) which migrate from the lateral ventricle into the olfactory bulb (OB) via the rostral migratory stream (RMS, Lois & Alvarez-Buylla, 1994; Lois et al., 1996); NPC which migrate from the SGZ travel a much short distance into the dentate gyrus (DG, Rickmann et al., 1987; Seki et al., 2007). The rate at which migration occurs within these regions is dependent on environmental and physiological conditions. Aging greatly impacts neurogenesis within both regions, as animals get older there are decreases in the total number of proliferative cells in both the SVZ (Apostolopoulou et al., 2017; Enwere et al., 2004) and SGZ (Kuhn et al., 1996). These alterations in the substructure of the neurogenic regions have downstream effects that can observed within the RMS (Capilla-Gonzalez et al., 2013), OB (Enwere et al., 2004; Tropepe et al., 1997), and DG (Jin et al., 2003).

Migration of NPC within the brain of aged animals has primarily been observed using histological techniques, however, there is a growing literature using different imaging modalities to monitor endogenous NPC continuously in live animals (Velde et al., 2012). MRI paired with intraventricular injections of micron sized iron oxide particles (MPIO) allows for high resolution *in vivo* monitoring of endogenous NPC migration (Shapiro et al. 2006a). Cells labelled endogenously have been shown to progressively accumulate within the OB, with levels of cells depending on the location of injection and volume of MPIO (Vreys et al., 2010; Granot et al., 2011). The optimal technique labels approximately 30% of NPC within the RMS (Sumner et al., 2009) and within the OB, MPIOs were found primarily in immature neurons (Nieman et al., 2010). Monitoring NPC migration using MRI has been used to understand homing of these cells in animal models of several disease states, including ischemia (Yang et al., 2008; Granot & Shapiro, 2014; Zhang et al., 2016). Cells in these experiments migrate away from their original targets and toward the disease. The presence of these cells in ischemic regions is hypothesized to improve neurological outcomes by promoting angiogenesis and by functioning as neuroprotective agents (Marlier et al. 2015). However, stroke and other disease that recruit NPC are more prevalent in older individuals (Wang et al., 2013) and the neurogenic response of these cells to disease is related to the age of animals (Gao et al., 2009). No studies have thus far used MPIO-mediated cell tracking in aged animals.

In this study we utilize a well-established technique of endogenously labelling NPC within the SVZ using MPIO, to observe difference in migration of cells into the OB across age. Histological studies have demonstrated a static snapshot of the progressive deterioration in NPC migration away from the SVZ and SGZ. We too observe these changes as animal age, with lower DCX staining found along the RMS and the DG in older animals. We additionally demonstrate *in vivo* that older animals have slower migration rates from the SVZ across 12 days and lower total amounts of NPC infiltration within the OB at the completion of the experiment. We also for the first time, demonstrate preliminary data showing MPIO uptake within the DG and lower uptake in the region within older animals. Neurological diseases that benefit from treatment with NPC have spark an interest in the utilization of endogenous NPC, however, these diseases often occur within aged population. Understanding and developing methods to monitor endogenous NPC in older animals is critical to developing effective strategies to overcome the detrimental effects of age on NPC populations.

## Methods

### 2.1 Animals Procedures

Animals were maintained under standard conditions in terms of temperature/humidity and were housed in a 12:12 Light/dark cycle with food and water *ad libitium*. All procedures were approved by the Institutional Animal Care and Use Committee at Michigan State University and are in accordance with the National Institutes of Health Guide for the Care and Use of Laboratory Animals.

Twenty-three adult Fischer 344 rats (Charles River, Raleigh, NC) underwent stereotactic surgery to inject MPIOs into the lateral ventricle. Intraventricular injections endogenously label neuroprogenitor cells within the SVZ, which can then be monitored longitudinally as cells migrate along the RMS into the olfactory bulb using MRI (Shapiro et al, 2006a). Animals were separated into groups based on age: 9 weeks old (n=6), 4 months old (n=5), 9 months old (n=6), and 2 years old (n=4). Serial MRI scans monitored the movement of NPCs over the course of 12 days, after which animals were sacrificed via transcardial perfusion. Brains were then scanned *ex vivo* using a high-field 9.4T magnet at 50μm resolution and then processed for histology using immunofluorescence.

### 2.2 Intraventricular Injection of MPIO for *In Situ* NPC Labeling

Prior to imaging, all animals were given intraventricular injections of Flash Red Fluorescent 1.63 μm MPIOs (ME04F, Bangs Laboratories, Fishers, IN) as previously described (Shapiro et al, 2006; Granot et al., 2011). In brief, animals were anesthetized with 2.5% isoflurane via endotracheal intubation using a 14G 2” catheter (ETT-16–50, Braintree Scientific, Inc., MA). Animals then were secured into a stereotaxic and a 3mm burr hole drilled into the skull. A 50μL pressure injection was made using a Hamilton syringe with a 33G needle at AP: −0.3mm, ML: 1mm, DV:-3.5mm (Figure S1). The needle was allowed to rest for 10 min and then slowly removed over the span of an additional 10 min. The skull was closed with bone wax and sutured without autoclips, as animals were immediately scanned with an MRI.

### 2.3 *In vivo* MRI

Each rat was scanned 5 times over the course of 12 days: on day 0, immediately after injection, and then on days 1, 3, 6 and 12 post-injection. Prior to scanning, animals were intubated and secured onto the animal bed via ear and tooth bars. A 2×2 rat brain coil (Bruker Corporation, MA) was placed onto the head and animals were scanned with a 7T Bruker Biospec 70/30 USR using a T1 Flash sequence with the following parameters: TE = 10.44 ms, TR = 31 ms, FOV = 2.56 × 2.56 × 2 cm, resolution = 100 × 100 × 100 μm, Averages = 2 and a total scan time = 47 min. Body temperature and breathing were continuously monitored (Model 1025; SA Instruments, Inc, NY). Animals were then given 1 mL of warmed saline after the 47 min scan and allowed to fully recover before being returned to their home cage.

### 2.4 *Ex vivo* MRI

After the final *in vivo* scan, animals were transcardially perfused with a 4% Paraformaldehyde (PFA) solution doped with 1 mM Gd-DTPA (Magnevist, Bayer, NJ). Brains were extracted with extra care to preserve the olfactory bulbs and post-fixed for at least 24hr in the Gadolinium-doped PFA. Brains were then placed into a 15mL Falcon tube filled with PBS. Tubes were fit into a volume coil and scanned on a 9.4T Bruker AVANCE at high resolution using a T1 Flash sequence with the following parameters: TE=10 ms, TR=30 ms, FOV =2.56 × 1.92 × 1.92 cm, resolution = 50 × 50 × 50 μm, Averages=10 and a total scan time= 12 hr 17 min.

### 2.5 Image Analysis

The images acquired during the *in vivo* scans were analyzed with the FIJI (Schindelin et al., 2012) Measure tool to determine the distance travelled by the particles from the SVZ to the olfactory bulb over time. The total volume of the particle distribution within the RMS and olfactory bulbs were measured with the *ex vivo* scans using the FIJI Analyze Particle tool after Thresholding. 3D Slicer 4.8 (https://www.slicer.org/; Fedorov et al., 2012) was then used to create a 3D volume rendering of the RMS for a representative animal in each age group.

### 2.6 Immunofluorescence Histology

Brains were soaked for at least 24hr in 30% sucrose after the completion of imaging, then imbedded in OCT and cut using a cryostat (Leica, Frankfurt, Germany) into 30μm sagittal sections. Free floating sections were stained for doublecortin (DCX). First, sections were blocked in 2% normal goat serum for 1hr and then incubated for 24hr at 1:2,000 of the primary antibody, rabbit anti-DCX (ab18723, Abcam). Sections were washed three times in PBS (10 min/wash) and then transferred into the 1:500 secondary antibody solution, goat anti-rabbit (ab150077, Abcam), for 1hr. Sections were mounted onto gelatinized slides, allowed to dry for 10 min and immediately coverslipped with ProLong Diamond Antifade Mountant (P36961, Life Technologies). Slides were allowed to cure at room temperature overnight and then stored at -20^°^C until microscopy.

Gross images of the RMS were acquired at 10× magnification for all samples using a Leica DMI4000B inverted fluorescence microscope. These images were used to determine the thickness of the RMS at the medial bend of the stream (Corona et. al, 2016) with the FIJI software and the Measure tool. Images were additionally acquired at a higher magnification (20×) using a Nikon A1 Rsi confocal laser scanning microscope at three locations along the RMS: (1) posterior near the SVZ, (2) medial bend, and (3) anterior near the olfactory bulb.

### 2.7 Statistical Analysis

The statistical software SPSS (IBM, Armonk, NY) was used for all analysis. One-way ANOVAs were used to compare age groups. Groups were compared to one another for significant ANOVAs using independent samples t-test. Significance was defined as *p*<0.05.

## Results

### 3.1 MRI demonstrates the impact of age on the rate of *in vivo* NPC migration

Many histological studies have observed the impact of age on the presence of NPC within the RMS, examining the number cells and size of the structure. Here, we examine for the first time the *in vivo* effects of age on NPC movement from the SVZ to the OB across 12 days (Figure 1). Iron oxide particles injected into the rostral horn of the lateral ventricle can be observed immediately post injection (1hr) as a dark blooming artifact within the structure. Within 24hrs, MPIOs are taken up by neural stem cells within the SVZ and can be observed beginning their journey along the RMS in the 9 wk, 4 mo, and 9 mo animals, heading down from the SVZ and then arcing upward toward the OB. No labelling or migration is observed in 2 year old animals until day 12. Younger animals (8 wk and 4 mo) have NPC that reach the OB by the third day, with more prominent and further migration noted in the youngest 8 wks cohort by day 12.

**Figure 1.**
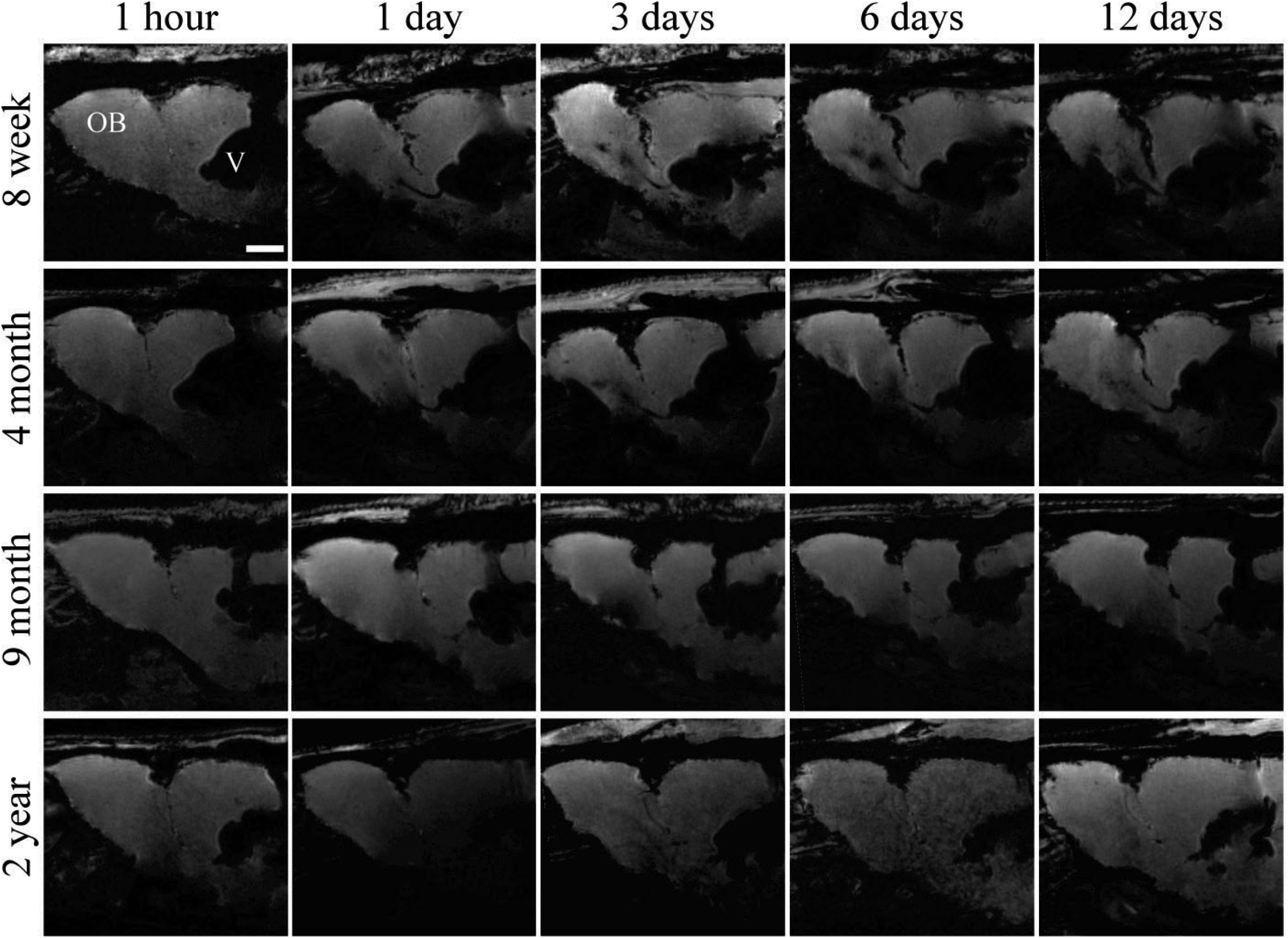
*In vivo* 7T MR images of representative animals from each age group at five time points post-injection of MPIOs. The RMS (or lack thereof) can be observed extending from the lateral ventricle (V) into the olfactory bulb (OB). Scale bar in the top left image represent 2000 µm.

Quantification of the migration using FIJI, verified the trends observed within the images obtained from the *in vivo* scans (Figure 2). There were significant main effects of age (F(3,15) = 43.1, *p*<0.001) and time (F(1,15) = 289.9, *p*<0.001), but also an interaction between age and time (F(3,15) = 25.9, *p*<0.001). Generally, there was more migration as time progressed for all groups, however, the pattern of migration over time was different between the age groups. When the age groups were statistically compared at each time point amongst one another, after 24hrs all the ages group were significant different from every other group (Figure 2). The younger 9 wk and 4 mo groups had similar non-linear curve over time, but the 4 mo group had significantly lower values after the initial 1hr scan demonstrating that the NPC of the 4 mo group covered less distance over time than the 8 wk animals. In the older animals there was a more pronounced difference in the curves observed compared to the younger animals, within the first 6 days the 8 mo group had a linear curve with depressed distance traveled by the NPC and the 2 yr group had no migration. The rate of migration for these animals was, therefore, much lower than in the younger groups (Table 1). Demonstrating that as animals age, the total distance traveled and rate of migration decreased.

**Table 1.** Rate of NPC migration (µm/h) across age.

|  | ( 0-1 ) | ( 1-3 ) | ( 3-6 ) | ( 6-12 ) |
| --- | --- | --- | --- | --- |
| **8 wk** | 100.8±3.4 | 31.5±4.4 | 15.7±3.4 | 6.0±1.8 |
| **4 mo** | 70.5±2.9 | 25.8±3.8 | 10.9±1.9 | 5.2±0.8 |
| **8 mo** | 9.4±9.4 | 13.6±6.5 | 14.7±4.0 | 6.8±1.0 |
| **2 yr** | 0 | 0 | 0 | 5.0±0.2 |

Note: Reported as mean ± SEM.

**Figure 2.**
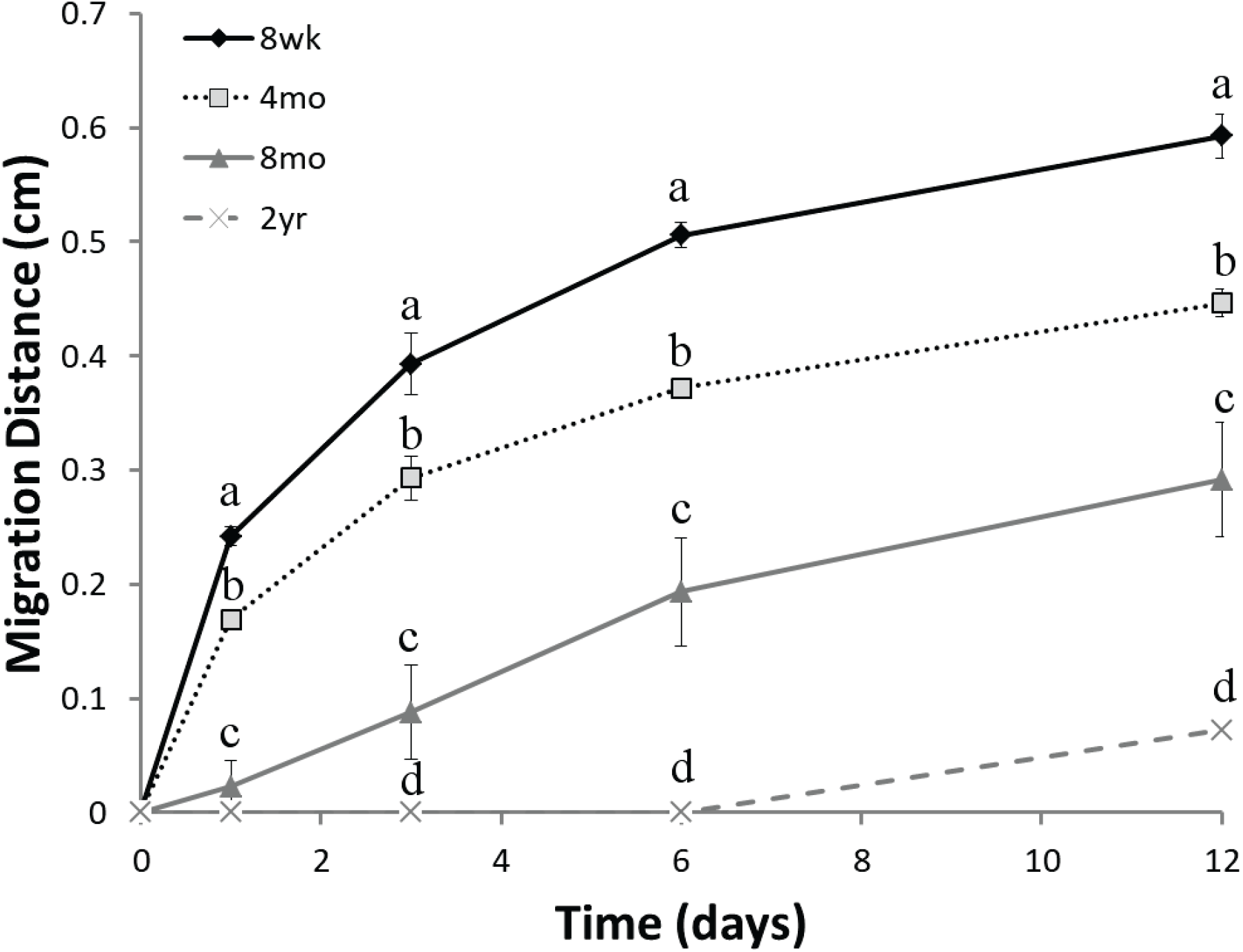
Quantitative evaluation of migration distance for all rats in each age group across time. Points represent mean of each age group at each time point ±SEM. Different letters represent significance difference (*p*<0.05) between age groups at each time point.

### 3.2 High Resolution imaging reveals the distribution of MPIO labelled cells within the OB across age

To identify the level of penetration into the OB by cells from the SVZ, we examined high resolution coronal sections and quantified the percent labelling of migrating cells between ages (Figure 3A-D). There was a significant effect of age in the total percentage of hypo-intensity seen within the interior of the OB (F(3,17) = 11.145, *p*<0.001). T-tests between the groups showed a significant difference between all ages for the number of cell migrating to the olfactory bulb (Figure 3E). When we examined the passive uptake of particles by quantifying the percentage of hypo-intensity in the outer portion of the OB, we found no significant difference between the groups (F(3,17) = 1.031, *p* = 0.409; Figure 3F). This demonstrates that across all age groups the same level of particles is taken up by cells that line surfaces exposed to CSF which contains MPIO.

**Figure 3.**
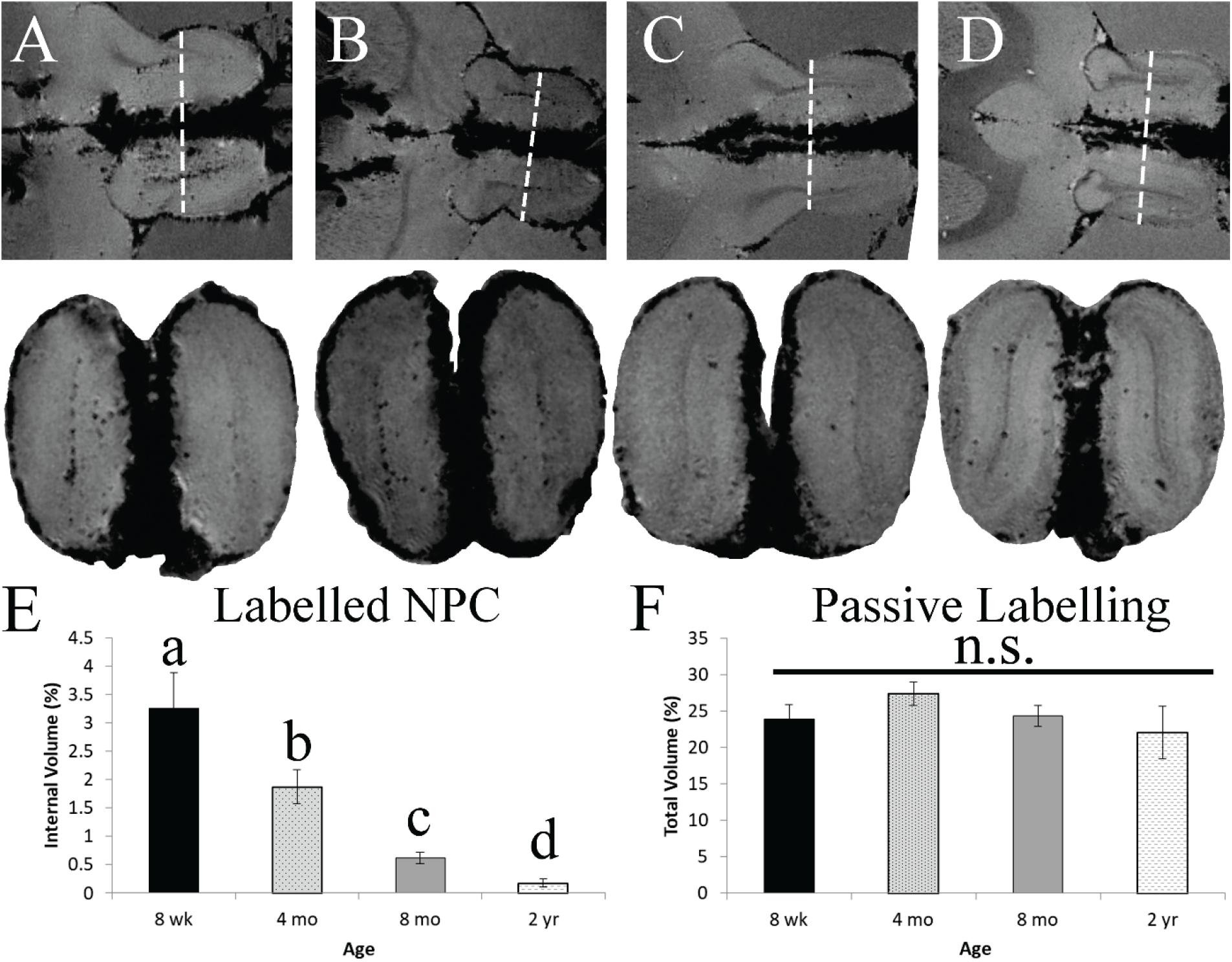
*Ex vivo* MR images in horizontal and coronal planes and analysis of MPIO quantity within the olfactory bulb. (A-D) Demonstrate the horizontal view (upper panel) of the RMS with a line indicating the level of the coronal section (lower panel). (E) Quantifies the percentage of hypointense area found in the internal portion of the coronal section at each age group. The different letters indicate groups that are significantly different from one another (*p*<0.05). (F) Quantifies the percentage of total hypointense area found outside of the internal labeling, which is cause by uptake of particles from the CSF by non-migrating cells.

To better demonstrate the overall labelling of NPC within animals, we generated 3D volume rendering for a representative animal in each age group (Figure 4). Visualizing the RMS in 2D slices does not provide a full view of all migrating cells, this was particularly evident in older animals who had fewer labeled cells. These renderings, however, verify what was observed in the 2D analysis with the total distribution of cells greatest in the youngest animal and the least labelling in the oldest.

**Figure 4.**
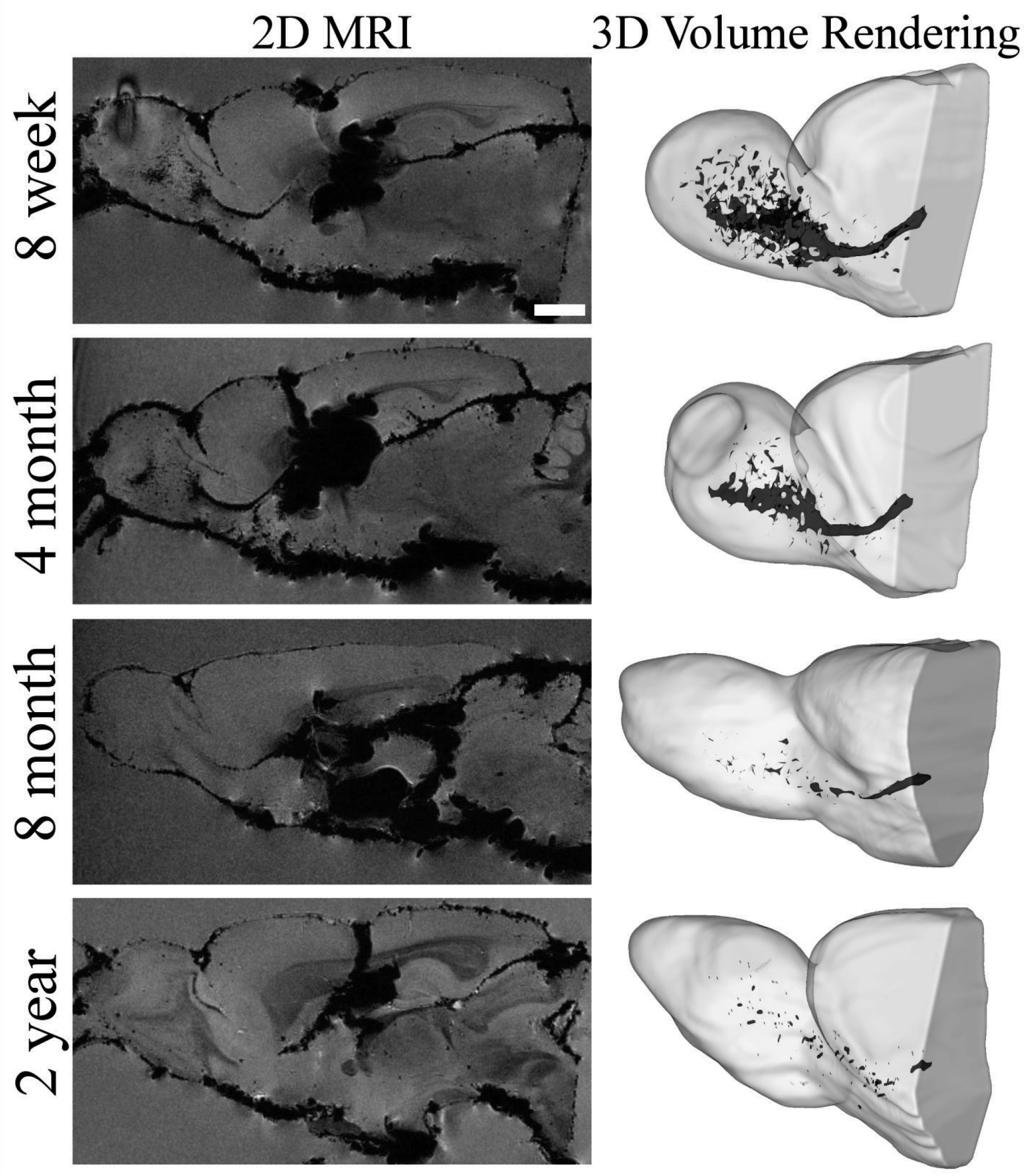
*Ex vivo 9.4T* MR images of the RMS displayed in two and three dimensional format for all age groups. The left column has examples of the best representative slices of the RMS in 2D, while the right column is a 3D rendering of the same animal with all MPIO containing cells of the RMS highlighted in dark grey. Scale bar in the top left image represent 2000 µm for the 2D images.

### 3.3 Histological comparison of DCX-labelled cells within the RMS mirrors MRI findings

To verify the differences seen with MRI of age, we additionally examined the histology of brain section using DCX labelling (Figure 5A), a marker associated with NPC. Using images attained with a fluorescent microscope, we measured the width of the RMS at the medial bend, were the stream is widest. There was a significant difference in width between the age groups (F(3,15) = 69.340, *p*<0.001). T-tests between the groups showed a significant difference between all ages, with smaller lengths as animals aged (Figure 5B). Confocal microscopy of three regions across the RMS demonstrated that at all levels of the structure, older animals had less DCX labelling for all regions photographed (Figure 6).

**Figure 5.**
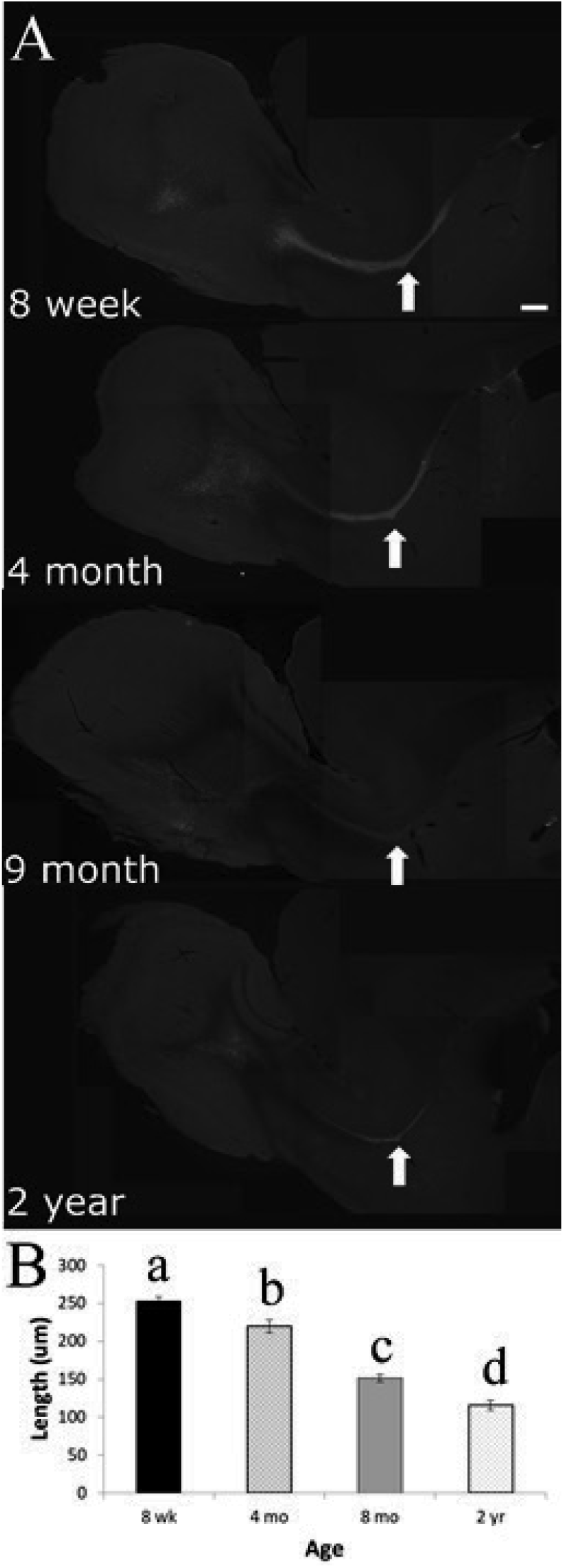
Fluorescent microscopy images and analysis of DCX labeling within the brains of animals across all age groups. (A) Representative images for the four age groups, the arrow denotes the medial bend of the RMS where measurements of length were taken. The scale bar represents 500 µm. (B) Graphical representation depicting the mean ±SEM of thickness (length, µm) of the RMS at the medial bend for all age groups. Different letters denote significant difference between the groups (*p*<0.05).

**Figure 6.**
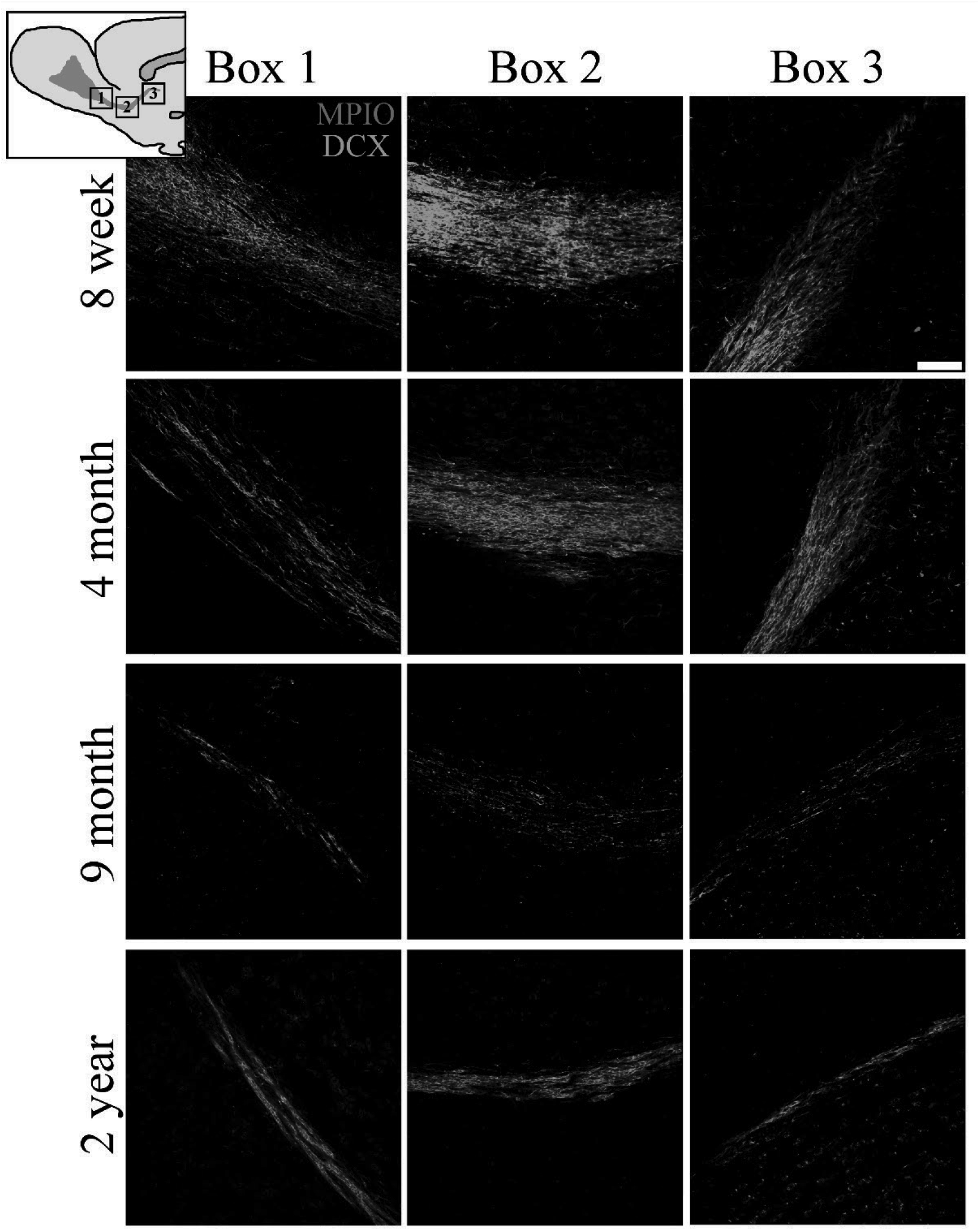
Confocal microscopy images of three locations along the rostral migratory stream for all age groups. The diagram in the upper left corner denotes the location of the images taken along the RMS: Box 1 is positioned anterior near the olfactory bulb, Box 2 at the medial bend of the RMS, and Box 3 is posterior near the SVZ. A representative from each age was selected for each row. The green within each image is DCX staining and the red represent fluorescent MPIOs. The scale bar on the top right image is 100 µm.

### 3.4 Novel identification of age-dependent MPIO labelling within the DG of the hippocampus

The DG of the hippocampus also has a population of neuroprogenitor cells, this region however has never been examined for labeling with MPIOs. Here we make a novel discovery of age-based variation in MPIO labelling within the DG. In our *in vivo* scans blooming artifacts caused by the presence of MPIOs within the ventricle obscure the visibility of the hippocampus (Figure S2). Post perfusion these particles were reduced around the structure, allowing for high resolution imaging of the region. *Ex vivo* scans within the caudal portion of the DG showed a marked uptake of particles within young animals and a decrease in uptake as animals aged (Figure 7A). Histological labeling of the rostral DG demonstrates that a younger animal (Figure 7B) has greater DCX labelling and more MPIOs in the region than an older animal (Figure 7C). These findings are preliminary, but suggest that the DG could also be labeled with MPIO and provide an avenue to monitor NPC within the hippocampus if the cells can be shown to definitively uptake MPIOs.

**Figure 7.**
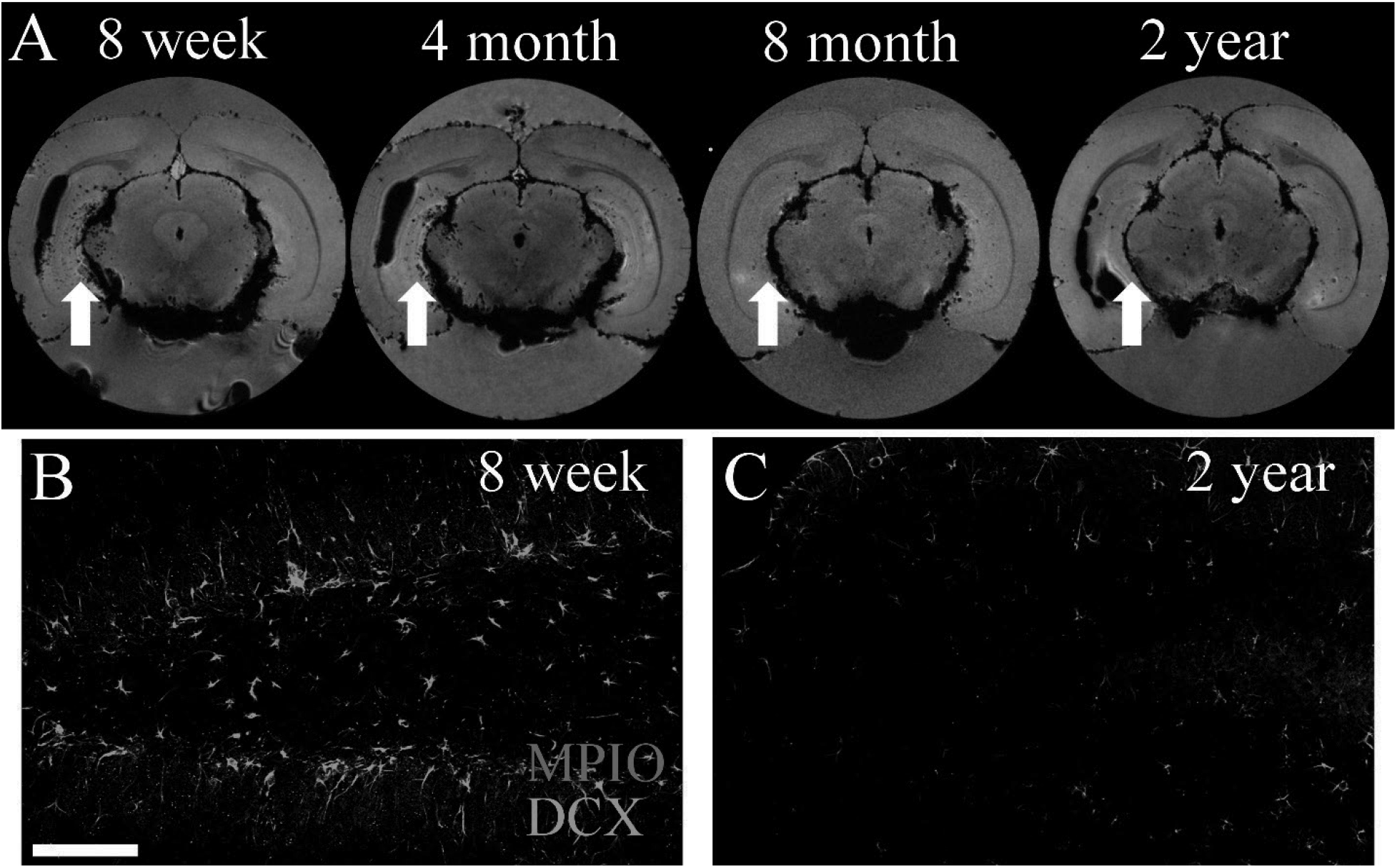
MPIO labeling within the dentate gyrus of the hypothalamus. (A) Representative 9.4T MR images acquired in *ex vivo* samples, dark hypotensive spots represent cells labeled with MPIOs. The white arrows denote the dentate gyrus in animals at each age. (B) Confocal images of a young 8 week old rat, DCX cells (green) and MPIO particles (red). (C) Confocal images of a 2 year old rat with little labeling of either DCX or MPIOs within the DG.

## Discussion

In this study, we demonstrate the age-based differences in NPC migration from the SVZ to the OB *in vivo*, verify the findings with high resolution *ex vivo* MRI and histology. We also highlight the possible use of the technique to observe NPC migration within another neurogenic region, the dentate gyrus. Endogenously labeling NPCs with iron oxide particles is a well-established technique for monitoring *in vivo* the migratory behavior of NPC cells from the SVZ into the OB using MRI (Shapiro et al., 2006a). This study implements the MPIO-mediated cell tracking to demonstrate the continued decrease in olfactory neurogenesis *in vivo* across age, impacting the rate of migration from the SVZ and the total cells successfully reaching the olfactory bulb. Understanding the physiology and behavior of these self-renewing cells, gives insight into how the brain responds and recovers from stressors, such as injury and disease

As rodents age, there is a clear impact on the neurogenic population of cells found in the subventricular zone. Histological studies have established that there are significant changes in the structure of the region, with a reduction in the total number of proliferative cells. Specifically, by middle-age animals have a significant reduction in the expression of Sox2 and DCX (Bouab et al., 2011) and by old age (+22 months) the cells are almost completely depleted (Capilla-Gonzalez et al., 2013). These ultrastructural alterations within the SVZ lead to a dramatic change in the appearance of the rostral migratory stream, as 2 year old animals have almost no DCX-stained (Capilla-Gonzalez et al., 2013) or Brd-U-stained (Mobley et al., 2013) migrating NPC when compared to young controls. In our staining, we observed similar decreases across ages in the labeling of the RMS using DCX. Young, 8 week and 4 month old animals, had very strong labelling while older 9 month animals had weaker labeling and 2 year old animals had RMS that appeared significantly thinned (Figure 6). Our study further demonstrate that differences caused by age observed in histology are similar to those observed on MRI, *ex vivo* scans showed a decrease in the number of NPC labelled in the OB as animals aged (Figure 3).

Our results also demonstrated how age directly impacts the rate at which NPC migrate into the olfactory bulb (Figure 1 & 2) by observing the movement of cells *in vivo* using MRI. *In vivo* imaging though MPIOs-labelling and MRI allows for continued monitoring of NPC across time and within subjects. The technique was first established to monitor single cells in vitro (Dodd et al, 1999; Hinds et al., 2003; Foster-Gareau et al., 2003) and then applied to migrating exogenously-labeled cells *in vivo* (Shapiro et al., 2004; 2006b). Endogenously labeling cells within the SVZ was established by Shapiro et al (2006a), they showed that intraventricular injections of MPIOs labeled NPC within the SVZ which could be observed migrating into the OB. Since the initial discovery, there have been many studies establishing the technique suggesting the method for scientific applications (Sumner et al., 2009; Nieman et al., 2009; Pothayee et al., 2017) and for preclinical work (Yang et al., 2008; Granot & Shapiro, 2014; Guglielmetti et al., 2014). The rates of migration, 102–120µm/h on day 0–1, established in these original methodological studies (Nieman et al., 2009; Pothayee et al., 2017) are similar to those describe in this manuscript for young 8 wk animals (Table 1). Clearly, our data also shows a stark decrease in the migration speed observed in older animals. These findings bring into question the validity of using young animals for regenerative medicine studies, if the target disease population is old and therefore have different NPC levels and migratory capabilities.

The dentate gyrus of the hippocampus is another neurogenic population of cells negatively impacted by the aging process (Ben Abdallah et al., 2010). With steady decreases in the number of Nestin-positive cells, a marker for NPCs, observed as animals age from three weeks to twenty-four months (Encinas et al., 2011). In our *ex vivo* scans, we observed clear differences in the number and distribution of labeled cells within the dentate gyrus (Figure 5). In the 8 week old rats, there are many hypo-intensive dark spots associated with labeled cells distributed widely across the caudal portion of the hippocampus. The number of spots decreases in a similar fashion as the histological studies across age. Our preliminary histology, also demonstrates a decrease in the amount of DCX staining observed within the region between the youngest and oldest animals. However, in depth analysis is required to verify the which cells types uptake particles and to quantify the degree of labelling. To our knowledge, no one has reported the labeling of NPC within the dentate gyrus using MPIOs (however, see Pohland et al., 2017) nor have they shown an age based difference in the distribution of cells within the region using MRI.

## Conclusion

The use of MPIOs to monitor NPC migration gives a new dimension into the study of age-based neurogenesis within the SVZ and DG, allowing for the continual *in vivo* data collection across time within a single animal. The technique increases the toolbox for understanding the physiology of the cell population and, also, demonstrates the limitations that the region has for translational applications.

---

**Figure S1.**
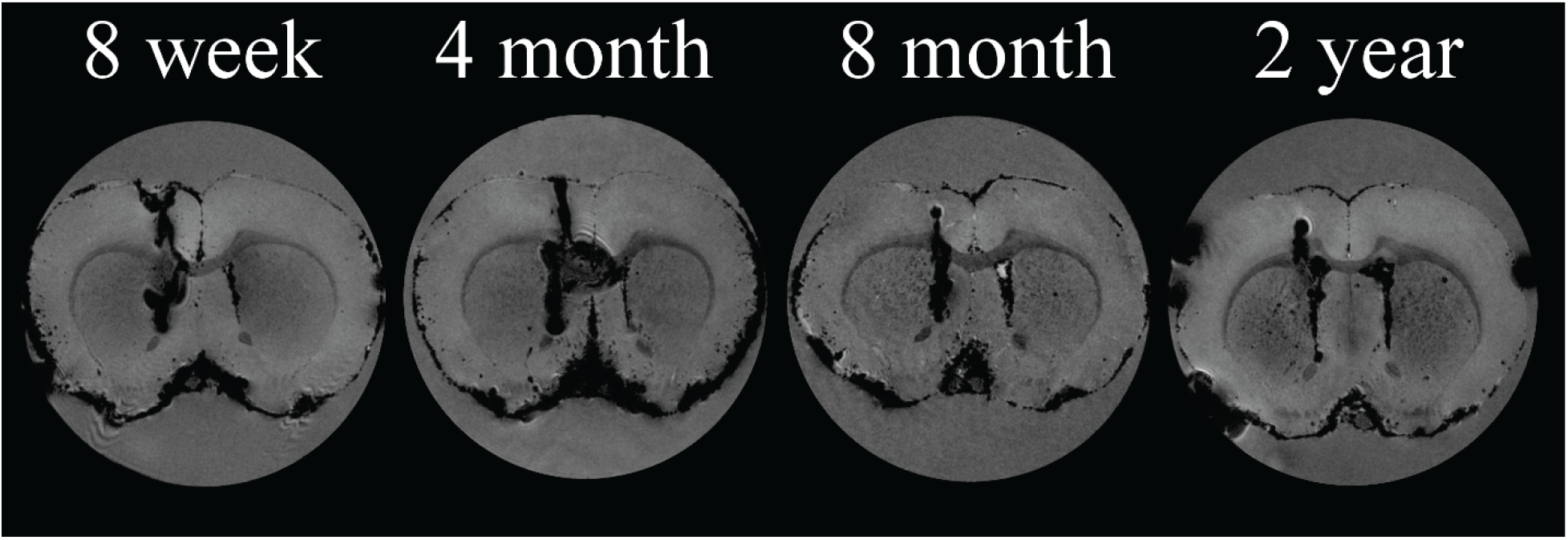
*Ex vivo* 9.4T MR images of representative animals from each age group at the level of intraventricular particle injection. The dark region passing through the cortex and corpus collosum and into the lateral ventricle is the needle track mark. This section demonstrates the consistency between age group of the stereotactic coordinates.

## References

Altman J (1969) autoradiographic and histological studies of postnatal neurogenesis. IV. Cell proliferation and migration in the anterior forebrain, with special reference to persisting neurogenesis in the olfactory bulb. J Comp Neurol, 137(4):433–457.

Apostolopoulou M, Kiehl TR, Winter M, de la Hoz EC, Boles NC, Bjorsson CS, Zuloaga KL, Goderie SK, Wang Y, Cohen AR, and Temple S (2017) Non-montonic changes in progenitor cell behavior and gene expression during aging of the adult V-SVZ neural stem cell niche. Stem Cell Reports, 9:1931–1947.

Capilla-Gonalez V, Cebrian-Silla, A, Guerrero-Cazares H, Garcia-Verdugo JM, and Quinones-Hinojosa A (2013) The generation of Oligodendroglial cells is preserved in the rostral migratory stream during aging. Front Cell Neurosci, 7:147.

Corona R, Retana-Marquez S, Portillo W, and Paredes RG (2016) Sexual behavior increases cell proliferation in the rostral migratory stream and promotes the differentiation of the new cells into neurons in the accessory olfactory bulb of female rats. Front. Neurosci,10:48.

Enwere E, Shingo T, Gregg C, Fujikawa H, Ohta S, and Weiss S (2004) Aging results in reduced epidermal growth factor receptor signaling, diminished olfactory neurogenesis, deficits in fine olfactory discrimination. The Journal of Neuroscience, 24(38):8354–8365.

Fedorov A, Beichel R, Kalpathy-Cramer J, Finet J, Fillion-Robin J-C, Pujol S, Bauer C, Jennings D, Fennessy FM, Sonka M, Buatti J, Aylward SR, Miller JV, Pieper S, and Kikinis R (2012) 3D Slicer as an Image Computing Platform for the Quantitative Imaging Network. Magn Reson Imaging, 30(9):1323–41. PMID: 22770690. PMCID: PMC3466397.

Gao P, Shen F, Gabriel RA, Law D, Yang E, Yang GY, Young WL, and Su H (2009) Attenuation of brain response to VEFG-mediated angiogenesis and neurogenesis in aged mice. Stroke 40(11):3596–3600.

Granot D, Scheinost D, Markakis EA, Papademetris X, and Shapiro EM (2011) Serial monitoring of endogenous neuroblast migration by cellular MRI. Neuromage, 57:817–824.

Granot D and Shapiro EM (2014) Accumulation of Micron Sized Iron Oxide Particles in Endothelin-1 Induced Focal Cortical Ischemia in rats is independent of cell migration. Magnetic Resonance in Medicine, 71:1568–1574.

Jin K, Sun Y, Xie L, Batteur S, Mao XO, Smelick C, Logvinova A, and Greenberg DA (2003) Neurogenesis and aging: FGF-2 and HB-EGF restore neurogenesis in hippocampus and subventricular zone of aged mice. Aging Cell, 2:175–183.

Kaplan MS and Bell DH (1984) Mitotic neuroblasts in the 9-day-old and 11-month-old rodent hippocampus. The Journal of Neuroscience, 4(6):1429–1441.

Kaplan MS and Hinds JW (1977) Neurogenesis in the adult rat: electron microscopic analysis of light radioautographs. Science, 197(4308):1092–1094.

Kuhn HG, Dickinson-Anson H, and Gage FH (1996) Neurogenesis in the dentate gyrus of the adult rat:age-related decrease of neuronal progenitor proliferation. The Journal of Neuroscience, 16(6):2027–2033.

Lois C, Garcia-Verdugo J, and Alvarez-Buylla A (1996) Chain migration of neuronal precursors. Science, 271(5251):978–981.

Lois C and Alvarez-Buylla (1994) Long-distance neuronal migration in the adult mammalian brain. Science, 264(5162):1145–1148.

Marlier Q, Verteneuil S, Vandenbosch R, and Malgrange B (2015) Mechanisms and functional significance of stroke-induced neurogenesis. Frontiers in Neuroscience, 9:458.

Nieman BJ, Shyu JY, Rodrigez JJ, Garcia AD, Joyner AL, and Turnbull DH (2009) In vivo MRI of neural cell migration dynamics in the mouse brain. NeuroImage 50:456–464.

Pohland M, Glumm R, Wiekhorst F, Kiwit J, and Glumm J (2017) Biocompatibility of very small superparamagnetic iron oxide nanoparticles in murine organotypoic hippocampal slice cultures and the role of microglia. International Journal of Nanomedicine, 12:1577–1591.

Rickmann M, Amaral DG, and Cowan WM (1987) Organization of radial glial cells during the development of the rat dentate gyrus. J Comp Neurol, 264(4):449–479.

Schindelin, J, Arganda-Carreras I, and Frise E et al. (2012) “Fiji: an open-source platform for biological-image analysis” Nature methods 9(7): 676–682, PMID 22743772, doi:10.1038/nmeth.2019

Seki T, Namba T, Mochizuki H, and Onodera M (2007) Clustering, migration, and neurite formation of neural progenitor cells in the adult rat hippocampus. J Comp Neurol, 502:275–290.

Shapiro EM, Gonzalez-Perez O, Garcia-Verdugo JM, Alverez-Buylla A, and Koretsky AP (2006a) Magnetic resonance imaging of the migration of neuronal precursors generated in the adult rodent brain. NeuroImage 32:1150–1157.

Sumner JP, Shapiro EM, Maric D, Conroy R, and Koretsky AP (2009) In vivo labeling of adult neural progenitors for MRI with micron sized particles of iron oxide: Quantification of labeled cell phenotype. NeuroImage 44:671–678.

Tropepe V, Craig CG, Morshead CM, and van der Kooy D (1997) Transforming growth factor-α null and senescent mice show decreased neural progenitor cell proliferation in the forebrain subependyma. The Journal of Neuroscience, 17(20):7850–7859.

Velde GV, Couillard-Despres S, Algner L, Himmelreich U, and van der Linden A (2012) In situ labeling and imaging of endogenous neural stem cell proliferation and migration. Wiley Interdiscip Rev Nanomed Nanobiotechnol, 4(6):663–679.

Vreys R, Velde GV, Krylychkina O, Vellema M, Verhoye M, Timmermans JP, Baekelandt V, and Van der Linden A (2010) MRI visualization of endogenous neural progenitor cell migration along the RMS in the adult mouse brain: Validation of various MPIO lebeling strategies. NeuroImage, 49:2094–2103.

Wang Y, Rudd AG, and Wolfe CDA (2013) Age and ethnic disparities in incidence of stroke over time: The South London Stroke Register. Stroke, 44:3298–3304.

Zhang F, Duan X, Lu L, Zhang X, Zhong X, Mao J, Chen M, and Shen J (2016) In vivo targeted MR imaging of endogenous neural stem cells in ischemic stroke. Molecules, 21:1143.

